# SeqEnhDL: sequence-based classification of cell type-specific enhancers using deep learning models

**DOI:** 10.1101/2020.05.13.093997

**Authors:** Yupeng Wang, Rosario B. Jaime-Lara, Abhrarup Roy, Ying Sun, Xinyue Liu, Paule V. Joseph

**Affiliations:** BDX Research & Consulting LLC, Herndon, Virginia, United States of America; Division of Intramural Research, National Institutes of Health, Bethesda, Maryland, United States of America

## Abstract

We propose SeqEnhDL, a deep learning framework for classifying cell type-specific enhancers based on sequence features. DNA sequences of “strong enhancer” chromatin states in nine cell types from the ENCODE project were retrieved to build and test enhancer classifiers. For any DNA sequence, sequential *k*-mer (*k*=5, 7, 9 and 11) fold changes relative to randomly selected non-coding sequences were used as features for deep learning models. Three deep learning models were implemented, including multi-layer perceptron (MLP), Convolutional Neural Network (CNN) and Recurrent Neural Network (RNN). All models in SeqEnhDL outperform state-of-the-art enhancer classifiers including gkm-SVM and DanQ, with regard to distinguishing cell type-specific enhancers from randomly selected non-coding sequences. Moreover, SeqEnhDL is able to directly discriminate enhancers from different cell types, which has not been achieved by other enhancer classifiers. Our analysis suggests that both enhancers and their tissue-specificity can be accurately identified according to their sequence features. SeqEnhDL is publicly available at https://github.com/wyp1125/SeqEnhDL.

## Introduction

Cell type-specific enhancers, *cis*-regulatory elements that up-regulate gene transcription in a cell type, play a key role in determining the regulatory landscape of the human genome (1). Enhancers are commonly located in the introns and immediately upstream of the transcription start site (TSS) of their target genes, but they are also known to populate gene deserts (2), reside in introns of neighboring genes (3) and co-localize with coding exons (4). Enhancer mutations are often associated with diseases. For example, mutations in a ∼400-bp sequence located 25 kb downstream of *PTF1A* are the most common cause of isolated pancreatic agenesis (5). A common sex-dependent mutation in an *RET* enhancer underlies Hirschsprung disease risk (6). Type 2 diabetes risk variants are enriched in islet enhancer clusters, some of which disrupt enhancer activity (7). While mutations within coding regions can be functionally distinguished between synonymous and non-synonymous mutations, functional inference of mutations in enhancers is challenging because little is known about enhancers’ sequence structures. Yet, accurate prediction of enhancers from DNA sequences is the basis of assessing whether mutation(s) could disrupt an enhancer’s activity, which is a type of mechanism for genetic diseases.

Evolutionary conserved regions (ECRs) were thought to be important components of enhancers, especially for developmental processes (8-10). ECRs were initially used for sequence-based enhancer prediction (11). Predicting enhancers based on transcription factor binding sites (TFBS) was proposed because TFBS tend to be conserved over vertebrate evolution (12-14). To ameliorate the uncertainty problem in conservation and TFBS information, direct sequence features such as *k*-mers were used to model enhancer prediction (15,16). These early studies did not achieve high prediction accuracy nor were they able to distinguish enhancers of different cell types.

With wide application of ChIP-seq technologies, enhancers were frequently profiled on a genome-wide scale (17). It is now known that enhancers are frequently associated with specific epigenetic marks such as H3K4me1 and H3K27ac (18,19). The ENCODE project produced genome-wide profiles of various epigenetic marks for multiple human cell types (20). By applying a hidden Markov model (i.e. ChromHMM) to these epigenetic marks, the sequence of the human genome has been binned into more than ten chromatin states, including enhancers (21,22). The “strong enhancer” state, shown to be associated with increased gene expression, provides genome-wide positioning of active enhancers for a cell type (21). Although the availability of these datasets renders positioning of enhancers unnecessary, the sequence structures of enhancers, especially their subtle differences among cell types, can be useful in understanding cell type-specific gene regulation and should be explored. Recent studies demonstrated that particular *k*-mers/TFBS are enriched in the epigenetic mark-determined enhancers, and a mutation might make an over-represented *k*-mer become insignificant or under-represented (23,24), which is highly consistent with the conservation theory.

In recent years, deep learning technologies have gained greater popularity than conventional machine learning methods. Due to the unprecedented effectiveness of deep learning technologies displayed in multiple applications, deep learning technologies have been adapted in biomedical research to address complex research questions (25). For example, Quang et al. (26) and Zhou et al. (27) used deep neural networks to annotate the effects of non-coding genetic variants. Alipanahi et al. (28) developed DeepBind to predict DNA- and RNA-binding proteins. Convolutional neural networks were adopted to predict enhancers (29-31). Quang and Xie (32) developed DanQ, a hybrid convolutional and recurrent deep neural network for quantifying the function of DNA sequences. Tan et al. (33) proposed an ensemble model of deep recurrent neural networks for identifying enhancers utilizing dinucleotide physicochemical properties.

Sequence-based enhancer identification has the potential of directly assessing the impacts of sequence changes on enhancers’ functionality (24,34). However, accurate identification of enhancers based on DNA sequences is challenging. The effectiveness of enhancer classifiers is influenced by proper generation of negative sequences. Negative sequences should contain similar basic sequence features with enhancers such as length distributions, GC and repeat contents (16,35,36); otherwise, enhancer classifiers may learn different nucleotide compositions rather than occurrences of key DNA motifs. Although there are many published studies regarding sequence-based enhancer prediction, it is still unknown whether these enhancer classifiers can distinguish enhancers from different cell types or tissues.

The sequence structures of enhancers may not be linear or additive. In fact, there could be complex grammar or semantics among different DNA elements that compose an enhancer (37,38). In this sense, deep learning models, which have shown unprecedented effectiveness in image, speech and language recognitions, should be more powerful in classifying enhancers. In this study, we propose SeqEnhDL, a deep learning framework for classification of cell type-specific enhancers based on sequence features. SeqEnhDL generates negative DNA sequences according to the GC contents of enhancers, and uses *k*-mer fold changes with a variety of *k*-mer lengths to build three deep learning models, including multi-layer perceptron (MLP), Convolutional Neural Network (CNN) and Recurrent Neural Network (RNN). The effectiveness and advantages of SeqEnhDL are demonstrated based on the chromatin state segmentation data of nine cell types from the ENCODE project (20).

## Methods

### Genome annotations

The sequences and transcripts of the human genome (hg19) were obtained from UCSC genome browser. The “knowngene” dataset was used to guide masking exons.

### Chromatin states

Chromatin state annotations of gm12878, H1hesc, hepg2, Hmec, Hsmm, Huvec, K562, Nhek and Nhlf cell types generated by ChromHMM (Broad version) were obtained from the ENCODE project. The data had a total of 15 chromatin states including 1_Active_Promoter, 2_Weak_Promoter, 3_Poised_Promoter, 4_Strong_Enhancer, 5_Strong_Enhancer, 6_Weak_Enhancer, 7_Weak_Enhancer, 8_Insulator, 9_Txn_Transition, 10_Txn_Elongation, 11_Weak_Txn, 12_Repressed, 13_Heterochrom/lo, 14_Repetitive/CNV and 15_Repititive/CNV. 4_Strong_Enhancer and 5_Strong_Enhancer states were used as enhancers in this study.

### Feature extraction

We masked exons and repetitive sequences of the human genome prior to retrieving DNA sequences for building enhancer classifiers. Initially, enhancers could have different lengths in folds of 200bp bins. We divided all initial enhancers into 200bp enhancer units and treated each enhancer unit as an enhancer. If any nucleotide of an enhancer was masked, the enhancer was removed from the analysis. The number of enhancers in each cell type is shown in Table 1. Each enhancer was a positive sequence. Both control and negative sequences were generated according to the GC contents of positive sequences. Control sequences, with a size of 3 folds of the positive sequences, were used to compute the background distributions of *k*-mers. Negative sequences, with the same size of the positive sequence set, were used as the negative set for training and testing enhancer classifiers. Fold changes of *k*-mer (k=5,7,9 and 11) frequencies (a pseudo count of 1 was added to both the denominator and numerator) were computed between the positive set and the control set and were used as feature dictionaries. Then, each 200bp sequence in the positive and negative sets was coded using the fold change of *k*-mers at each nucleotide position. Thus, for the CNN model, and the features for each sequence were a 200×4×1 array of *k*-mer fold changes. For the RNN model, the features for each sequence were a 200×4 array of *k*-mer fold changes. For MLP and conventional machine learning models, we flattened the features of each sequence into a vector of 800 *k*-mer fold changes.

**Table 1.** Number of enhancers in each cell type after the filtering procedure.

| **Cell type** | **Number of enhancers** |
| --- | --- |
| gm12878 | 92,763 |
| H1hesc | 21,560 |
| hepg2 | 69,337 |
| Hmec | 136,641 |
| Hsmm | 145,952 |
| Huvec | 147,113 |
| K562 | 80,912 |
| Nhek | 136,803 |
| Nhlf | 111,889 |

### Construction of machine learning models

Positive sequences for any model were initially split into training and testing sets according to 80:20 rule. Negative sequences were split according to the divisions of their corresponding positive sequences. For deep learning models, the initial training set was further divided into 70% training and 30% validation sets. Deep learning models were built based on the training and validation sets, and performance was assessed based on the testing set. Accuracies were defined as the proportion of correct classifications of the testing sequences. AUCs were computed based on the prediction scores on the testing sequences. Five-fold cross-validation was employed to generate reliable estimates of accuracies and AUCs (average from five runs).

### Deep learning models

Deep learning models were generated using the Tensorflow and Keras software, available from Python3.7. Parameters were chosen heuristically and consistently to reach fair comparisons among different machine learning models and different cell types. Parameters included batch size: 512; learning rate: 0.001; epochs: 20; optimizer: Adam; loss: categorical_crossentropy. The best model during the 20 epochs was saved and used for prediction on the testing data. For RNN, bidirectional long short-term memory (LSTM) was adopted. Structures of deep learning models are displayed in supplementary Table S1.

### Machine learning models involved in comparison

gkm-SVM was executed using default parameters. We also tested LS-GKM (39), which is the version of gkm-SVM for large datasets, and found the two software generated very consistent outcomes. Fasta sequences, rather than *k*-mer features, were fed into gkm-SVM. Due to high computational burden of gkm-SVM, 2000 positive sequences and 2000 negative sequences were randomly selected from the original training datasets to train enhancer classifiers, while for testing the entire dataset was used.

DanQ was executed using default parameters. Fasta sequences, which were extended to 1000bp from the 200bp bins, were fed to DanQ. DanQ made predictions on 919 ChIP/DNase-seq marks. For each cell type, the ChIP/DNase-seq mark with the highest accuracy was used to represent DanQ’s performance.

For Tan et al.’s enhancer classifier (33), original model structures and weights were downloaded and executed to classify the testing datasets of this study.

SVM models (linear and RBF kernels) and other conventional machine learning models including Decision Tree, Random Forest, AdaBoost, and Naïve Bayes were carried out using the scikit-learn python package. To ensure that computation was completed within one hour, 2000 positive sequences and negative sequences from the original training datasets were randomly selected for building conventional machine learning models. We found a sample size of 2000 sequences was enough to generate reliable and consistent estimates of accuracies for these conventional machine learning models. Parameter settings of conventional machine learning models are shown in supplementary Table S2.

#### Computational resource

All programs of this study were executed on the NIH Biowulf linux cluster. Tensorflow, Keras and required Python libraries were pre-configured under Python 3.7 on Biowulf. Deep learning jobs were executed on GPU nodes.

## Results

### The SeqEnhDL framework

The SeqEnhDL framework is depicted in Figure 1. The framework started from “bed” files containing the chromosomal positions of a large number (e.g. >1000) of enhancers. Proper data preprocessing and feature extraction are of key importance. For instance, the DNA sequences of enhancers were retrieved from the human genome where exon and repetitive sequences were masked. Then these DNA sequences were divided into individual enhancers with a fixed length of 200bp. An arbitrary enhancer length of 200bp makes features more standardized and comparable, although some of the divided enhancers may not contain enough enhancers’ motifs. Enhancer sequences composed the positive sequences. Control sequences for computing *k*-mer fold changes, and negative sequences for testing enhancer classifiers, were randomly selected from the genome where exon, repetitive and enhancer sequences were masked, according to the GC contents of enhancer sequences. *K*-mer (*k*=5, 7, 9 and 11) fold changes between enhancer and control sequences were computed and used for generating the feature at each nucleotide position of any positive or negative sequence. We chose odd *k*-mers because fold changes of different *k*-mers can be aligned at their central nucleotide position. Moreover, many 11-mers still have substantial occurrences because tens of thousands of enhancers are used to build *k*-mer dictionaries. Any dataset for building an deep learning enhancer classifier should be devided into training, validation and testing data, in a cross-valiation mode. Three deep learning models-MLP, CNN and RNN were built. Multilayer perceptrons are fully connected networks. CNN takes advantage of the hierarchical pattern in data and assemble more complex patterns using smaller and simpler patterns. RNN makes use of sequential information among features. Particularly, bidirectional long short-term memory (LSTM) RNN which can learn long-term dependencies was adopted.

**Figure 1.**
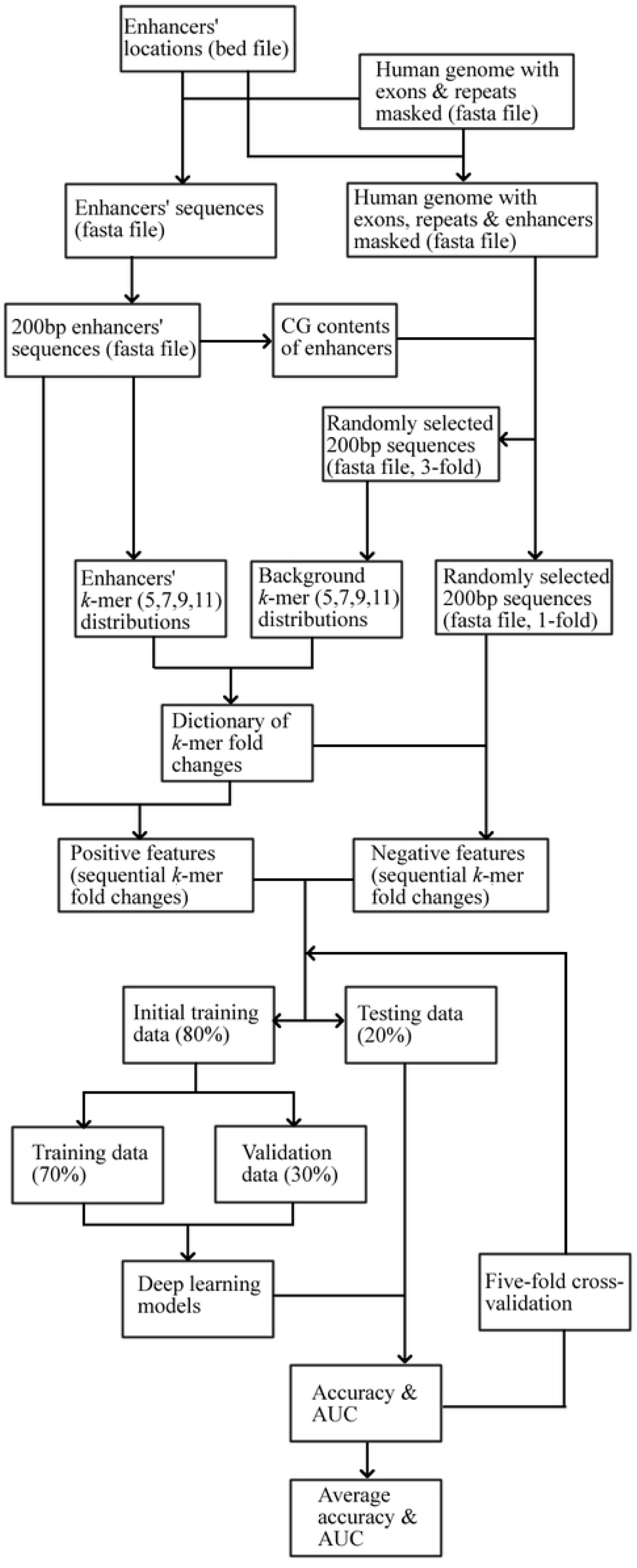
Flowchart of the SeqEnhDL framework.

### Evaluation of the performance of SeqEnhDL

Discriminating enhancers of a single cell type/tissue from randomly selected sequences have been studied before and provided the foundation for evaluating the performance of SeqEnhDL. We retrieved DNA sequences located within the “strong enhancers” chromatin states of nine cell types from the ENCODE project (20). Control sequences matching the GC content of each enhancer were constructed for each cell type (see Methods). The performance of SeqEnhDL was evaluated in terms of accuracy and area under the curve (AUC) for distinguishing enhancers in each cell type. State-of-the-art methods were selected for comparison with SeqEhnDL. gkm-SVM (16,35) was chosen for comparison because it uses *k*-mer information to predict enhancers. DanQ (32) was chosen for comparison because it is an RNN-based tool for predicting the functions of noncoding sequences. The performance of DanQ on each cell type was represented by the highest statistics among predictions on 919 ChIP/DNase-seq marks. When different tools were executed, five-fold cross-validation was employed in order to generate reliable performance measures. Comparisons of performances among different tools (Figure 2) show that all the three models of SeqEnhDL greatly outperform gkm-SVM and DanQ. The accuracies of SeqEnhDL range from 0.961 to 0.999, suggesting that enhancers can be accurately identified on different cell types. Comparison of Receiver Operating Characteristic (ROC) curves on the hepg2 cell type (Figure 3) re-conformed that SeqEnhDL performed better. Of note we also ran a recent approach based on ensemble of deep RNNs (33) for a comparison. However, its accuracies and AUCs were around 0.5 (supplementary Table S3), indicating that this compared approach was ineffective on the datasets of this study.

**Figure 2.**
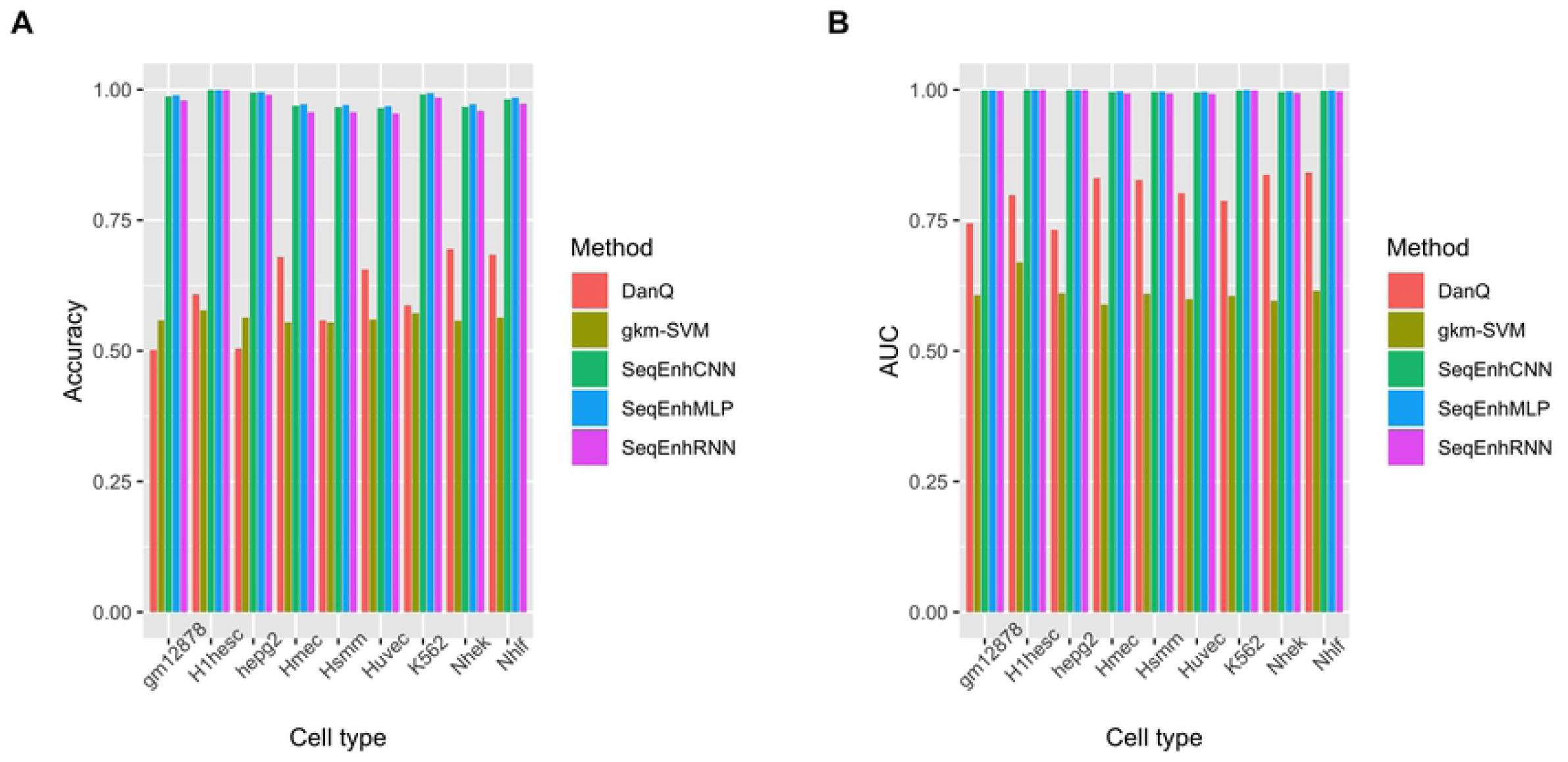
Comparison among different enhancer classifiers with regard to distinguishing cell type-specific enhancers from randomly selected non-coding sequences. (A) Comparison of accuracies. (B) Comparison of AUCs.

**Figure 3.**
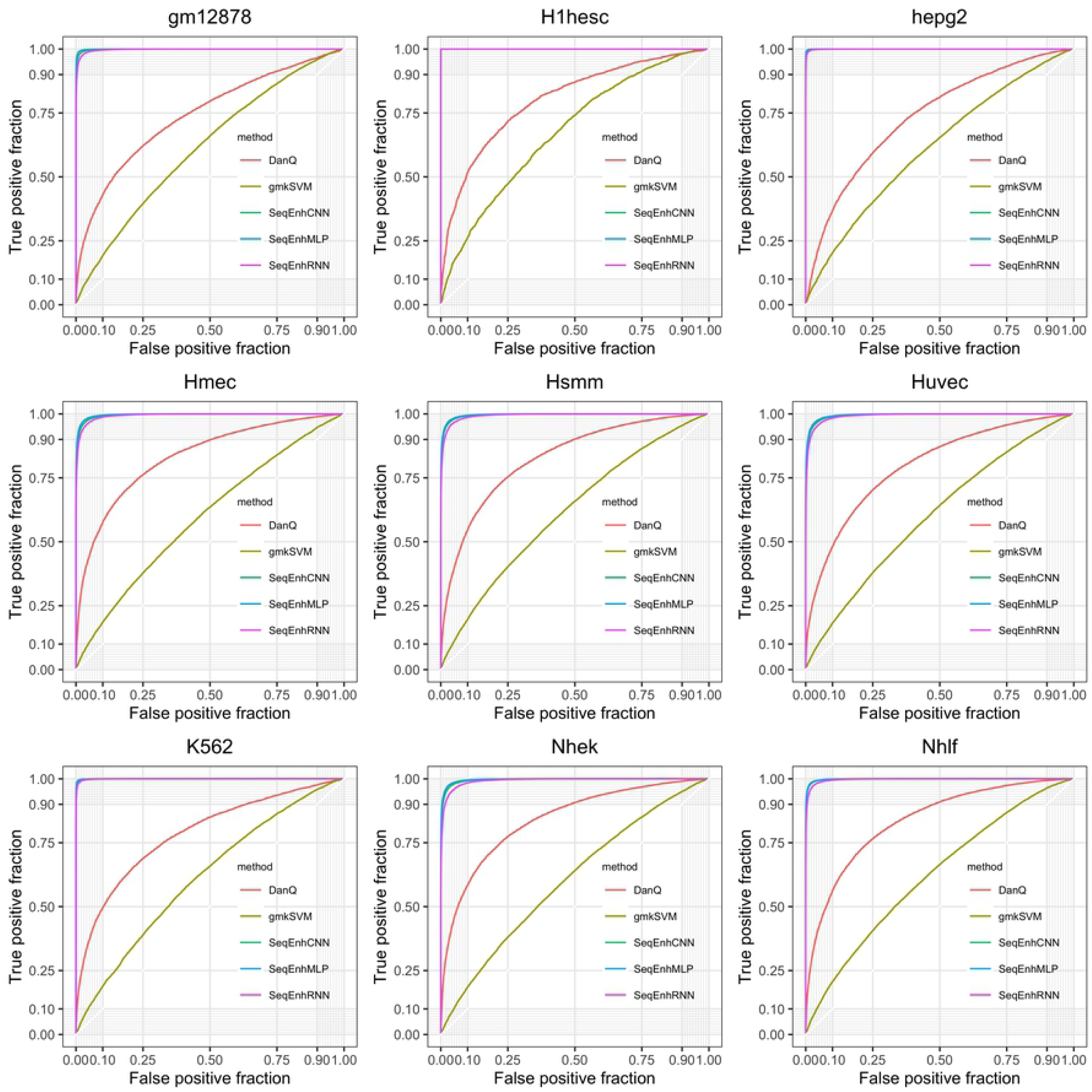
Comparison among different enhancer classifiers in terms of ROC curves. The curves were generated based on the first cross-validation dataset of each cell type.

Using *k*-mer features to predict tissue-specific enhancers has been utilized previously and achieved fair performance. For example, Lee et al. (15) developed a Support Vector Machine (SVM) framework based on *k*-mer features and achieved an AUC of 0.94 for EP300-bound enhancers. gkm-SVM (16) achieved an AUC of 0.947 for EP300-bound enhancers. However, the AUCs of gkm-SVM substantially decreased on the ENCODE datasets. In contrast with EP300-bound enhancers which tend to have binding sites for a limited number of transcription factors, enhancers determined by HMM models which are based on multiple tracks of epigenetic marks represent a more complete set of enhancers for the assayed cell type. Thus, the ENCODE enhancers have more complex DNA element structures and interdependencies. Therefore, the classification tasks of this study are more challenging.

To further demonstrate that deep learning models are better than conventional machine learning models based on the same features, we flattened the *k*-mer features and built enhancer classifiers based on six conventional machine learning models. Note that for each cell type 2000 postive and negative sequences were randomly selected and repeated 10 times in order to ensure training of each deep learning and conventional machine learning model could be finished within one hour. Figure 4 shows that accuracies of SeqEnhCNN and SeqEnhRNN are consistently higher than conventional machine learning models, and SeqEnhMLP is among the second tier in most cell types. These analyses collectively suggest that enhancers present in a single cell type can be accurately identified based on sequence features by SeqEnhDL, and SeqEnhDL greatly outperforms existing methods regarding discriminating enhancers from randomly selected sequences.

**Figure 4.**
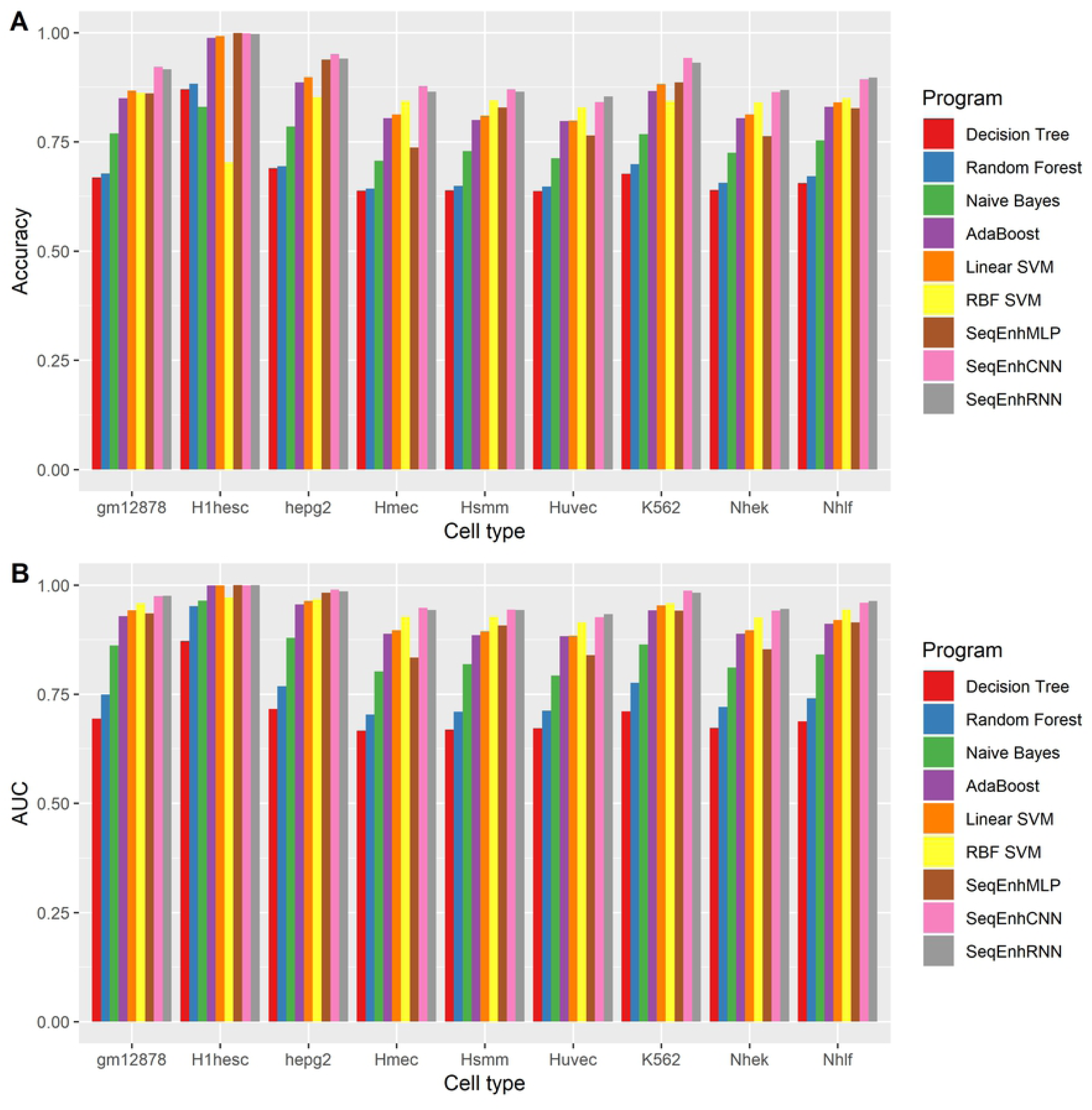
Comparison between SeqEnhDL and conventional machine learning models. (A) Comparison of accuracies. (B) Comparison of AUCs.

### SeqEnhDL can discriminate enhancers’ cell types based on DNA sequences

It is widely accepted that regulatory sequences tend to have different patterns of nucleotide usage relative to other types of DNA sequences, including sequence conservation and CG density (40). When enhancers from different cell types are discriminated, different classes of DNA sequences have similar basic sequence features. Successful machine learning models for distinguishing enhancers from different cell types must learn cell type-specific sequence structures such as domains, motifs and their interactions. Previous enhancer classifiers were not examined regarding this capacity.

Although most of previous enhancer classifiers were designed for distinguishing enhancers and non-enhancers, some may be adapted for distinguishing enhancers from different cell types by treating one cell type as the negative group. We applied gkm-SVM and SeqEnhDL to distinguish enhancers from different cell types. For each pair of cell types, we switched the assignments of positive and negative groups and computed the average accuracy and AUC. The accuracies and AUCs for all pairs of cell types are displayed in Figure 5. The accuracies and AUCs of gkm-SVM for all pairs of cell types are around 0.5, indicating that gkm-SVM failed to capture tissue-specificities of enhancers. In contrast, all models of SeqEnhDL generated high accuracies (e.g. >0.9) and AUCs (e.g. >0.95) in most cell type combinations, indicating that SeqEnhDL is able to identify tissue-specificity. This analysis suggests that SeqEnhDL can learn distinct sequence features related to tissue-specificities and discriminate enhancers from different cell types.

**Figure 5.**
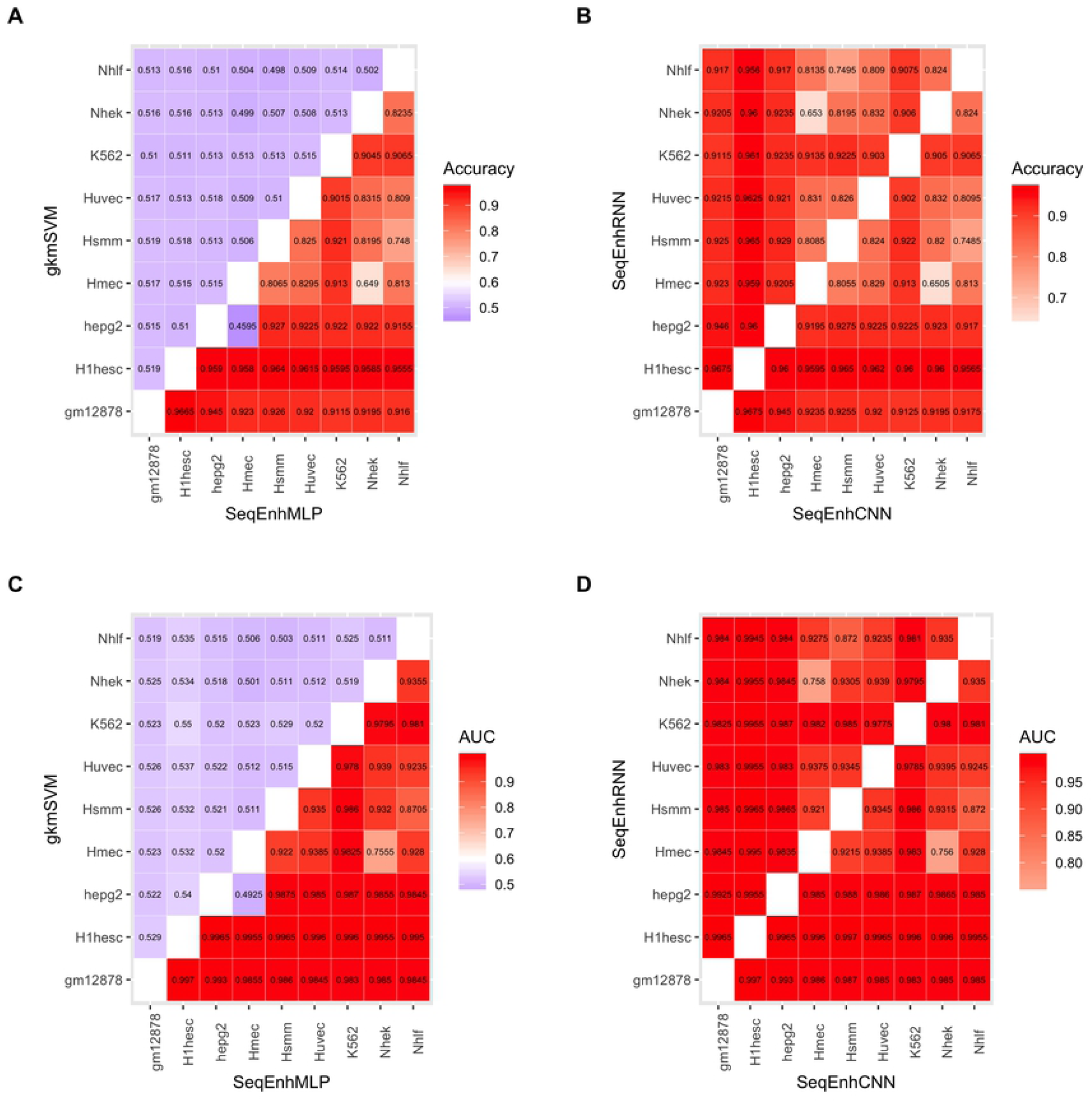
Comparison between gkm-SVM and SeqEnhDL with regard to discriminating enhancers from two cell types. (A) and (B) Comparison of accuracies. (C) and (D) Comparison of AUCs.

## Discussion

In this study, we propose SeqEnhDL, a feature extraction and deep learning framework for classifying cell type-specific enhancers based on sequence features. A variety of analyses were performed to demonstrate that SeqEnhDL outperforms existing enhancer classifiers. We further proved that SeqEnhDL can be used to discriminate enhancers from different cell types. Theoretically, when enhancers can be accurately distinguished, enhancer classifiers can be applied to predict whether mutations within enhancers may disrupt enhancer structures. Therefore, SeqEnhDL also provides a novel computational avenue for understanding the genetic mechanisms of disease-associated variants in non-coding regions.

Although previous studies for sequence-based enhancer classification achieved satisfactory performance scores (15,16), those models have limitations. First, some studies did not restrict control/negative sequences to noncoding sequences having similar basic sequence features with enhancers, making enhancer identification an easy problem and model performance greatly overestimated. Second, DNA motif-based methods relied on the frequencies of known transcription factor binding sites, leading to sequence features limited/biased to a priori-knowledge which could omit important unknown transcription factor binding sites. The SeqEnhDL framework has solved these limitations. Furthermore, utilization of CNN and RNN models have incorporated sequential dependencies among *k*-mers, thus leading to substantial improvement over existing approaches.

When we built enhancer classifiers, we separated control and negative sequences, which can significantly reduce the chances of overfitting, which is a common problem of machine learning models. We used all enhancers’ sequences to compute *k*-mer fold changes. Although theoretically enhancers can be divided into two subsets - one for computing *k*-mer fold changes and the other for testing enhancer classifiers, it is not practical because longer *k*-mers which are very important for composing enhancers may occur only few times in a cell type.

We succesfully applied SeqEnhDL to discriminate enhancers from two cell types. gkm-SVM failed to distinguish enhancers from different cell types, indicating that most (if not all) previous *k*-mer based models tend to learn the common features of enhancers rather than tissue-specific motif structures. This successful application suggests that tissue/cell type-specific gene regulation could be better understood based on machine learning of high-level enhancers’ structures. Continuing studies on multi-class enhancer classifiers will contribute to deeper understanding of the regulatory mechanisms of cell type-specific gene expression, and how non-coding mutations may cause disease phenotypes in different tissues (e.g. cancer from different tissues).

## Availability

Genome sequences and annotations were downloaded from UCSC genome browser (http://genome.ucsc.edu/). Chromatin state segmentation data were downloaded from the ENCODE project (http://hgdownload.soe.ucsc.edu/goldenPath/hg19/encodeDCC/wgEncodeBroadHmm/). Programs of this study were written in Perl, Python and R. All source code is freely available at https://github.com/wyp1125/SeqEnhDL. Testing datasets of SeqEnhDL for reproduction purpose are available at http://www.bdxconsult.com/SeqEnhDL.

## Supplementary data

Supplementary Data are available at PlosOne online.

## Acknowledgment

The authors would like to thank Dr. Joan Austin for her comments and editorial assistance.

## Funding

PVJ is supported by the National Institute of Nursing Research [1ZIANR000035-01]. PVJ is also supported by the Office of Workforce Diversity, National Institutes of Health and the Rockefeller University Heilbrunn Nurse Scholar Award[no award number]. RBJL received Intramural Research Training Awards, Office of Intramural Training & Education, National Institutes of Health, Department of Health and Human Services[no award number]. Funding for open access charge: National Institutes of Health.

## Conflict of interest

Potential competing interests: The authors have no potential conflicts of interest to disclose.

